# Hyperacetylated Chromatin Domains Mark Cell Type-Specific Genes and Suggest Distinct Modes of Enhancer Function

**DOI:** 10.1101/666784

**Authors:** Sierra Fox, Jacquelyn A. Myers, Christina Davidson, Michael Getman, Paul D. Kingsley, Nicholas Frankiewicz, Michael Bulger

## Abstract

Stratification of enhancers by relative signal strength in ChIP-seq assays has resulted in the establishment of super-enhancers as a widespread and useful tool for identifying cell type-specific, highly expressed genes and associated pathways. We have examined a distinct method of stratification that focuses on peak breadth, termed “hyperacetylated chromatin domains” (HCDs), which classifies broad regions exhibiting histone modifications associated with gene activation. We find that this analysis serves to identify genes that are both more highly expressed and more closely aligned to cell identity than super-enhancer analysis does when applied to multiple datasets. Moreover, genetic manipulations of selected gene loci suggest that at least some enhancers located within HCDs work at least in part via a distinct mechanism involving the modulation of covalent histone modifications across domains, and that this activity can be imported into a heterologous gene locus. In addition, such genetic dissection reveals that the super-enhancer concept can obscure important functions of constituent elements.

## Main

Enhancers are *cis*-regulatory DNA sequences that are bound by transcriptional activators which regulate genetically linked gene promoters^1–5^. They are also historically characterized by their ability to function over distances of anywhere from 100 bp to more than 1 Mb. Whole-genome methods of identifying enhancers rely on associated histone modifications, common transcriptional cofactors, and/or cell type-specific transcription factors (TFs). These methods suggest that mammalian genomes harbor a large number of enhancer sequences – perhaps 1-2 million – and that they represent the most numerous and significant *cis*-acting sequence determinants of cell type-specific gene expression.

Fundamental issues regarding enhancer function, however, remain unclear. For example, there is no consensus for how enhancers communicate with their cognate gene promoters. The dominant model, termed “looping,” involves direct interactions between factors bound to enhancers and factors bound near promoters. Evidence for such interactions, however, has provided little insight into how a distal enhancer finds a gene promoter, or how it distinguishes among potential promoters in gene-dense regions. Moreover, some evidence suggests that mechanisms of enhancer-promoter communication may be more varied^3,5^.

Additional insight into enhancer function can be obtained by discerning functional differences between them. One attempt to classify enhancers according to “strength,” as determined by signal intensity in genome-wide ChIP-seq assays for associated enhancer marks, has led to the classification of super-enhancers, which by various analyses have been shown to be associated with highly-expressed genes that define cell identity^6–8^. Obvious functional differences have not emerged from studies of super-enhancers, however, with the only reported distinction being a higher diversity and/or number of TF binding sites mapping to super-enhancers^9,10^. Mechanistically, there is as yet no indication that, aside from “strength” of activation, these sequences are intrinsically any different from other enhancers that exhibit weaker signal for enhancer-defining marks.

We have previously investigated enhancer-associated histone modifications from the perspective of peak breadth, as opposed to signal strength. We characterized specific regions as “hyperacetylated chromatin domains” (HCDs), defined as continuous genomic regions exhibiting significant enrichment for histone hyperacetylation and other marks associated with active transcription, most notably H3K4 dimethylation (H3K4Me2). From this analysis, we identified a novel enhancer within the murine β-globin locus, termed HS-E1, which is required for the formation of such a broad region of histone hyperacetylation encompassing the two genes within the cluster that are expressed during primitive erythropoiesis^11,12^. Our results suggested a distinction between enhancers that mediate broadly-distributed changes in chromatin structure vs. enhancers that work *via* other mechanisms.

To further investigate this, we have performed ChIP-seq analyses of specific histone modifications in primary murine erythroid, retinal, and intestinal epithelial cells and ranked peaks according to breadth. We find that an HCD ranking that utilizes a combination of two histone marks suffices to identify a subpopulation of genes that are both more highly expressed and more cell type-specific than those associated with super-enhancers. Moreover, deletion of enhancers found in loci with or without an HCD identifies functional differences between enhancers that have the ability to modulate long-range chromatin structure and those that do not. Insertion of an HCD-associated enhancer into another locus results in the formation of an HCD, further suggesting a distinct function for this class of enhancer.

## Results

Our prior studies suggested the existence of hyperacetylated chromatin domains (HCDs) controlled by enhancers at specific genomic regions, such as the murine β-globin locus^12^. In an effort to define the genome-wide occurrence and distribution of such domains, we performed and/or compiled a number of ChIP-seq and other datasets. These included (1) ChIP-seqs we performed using e14.5 murine fetal liver, which is comprised of 70-80% erythroid cells, using antibodies specific for H3K27 acetylation (H3K27Ac), and for H3K4 mono-, di- and trimethylation (H3K4Me1, H3K4Me2, H3K4Me3); (2) ATAC-seq^13^ we performed using sorted proerythroblasts from e14.5 murine fetal liver; and (3) publicly available ChIP-seq datasets from e14.5 murine fetal liver for the erythroid transcription factors GATA-1 and SCL/Tal1^14^ (Figure 1).

**Figure 1:**
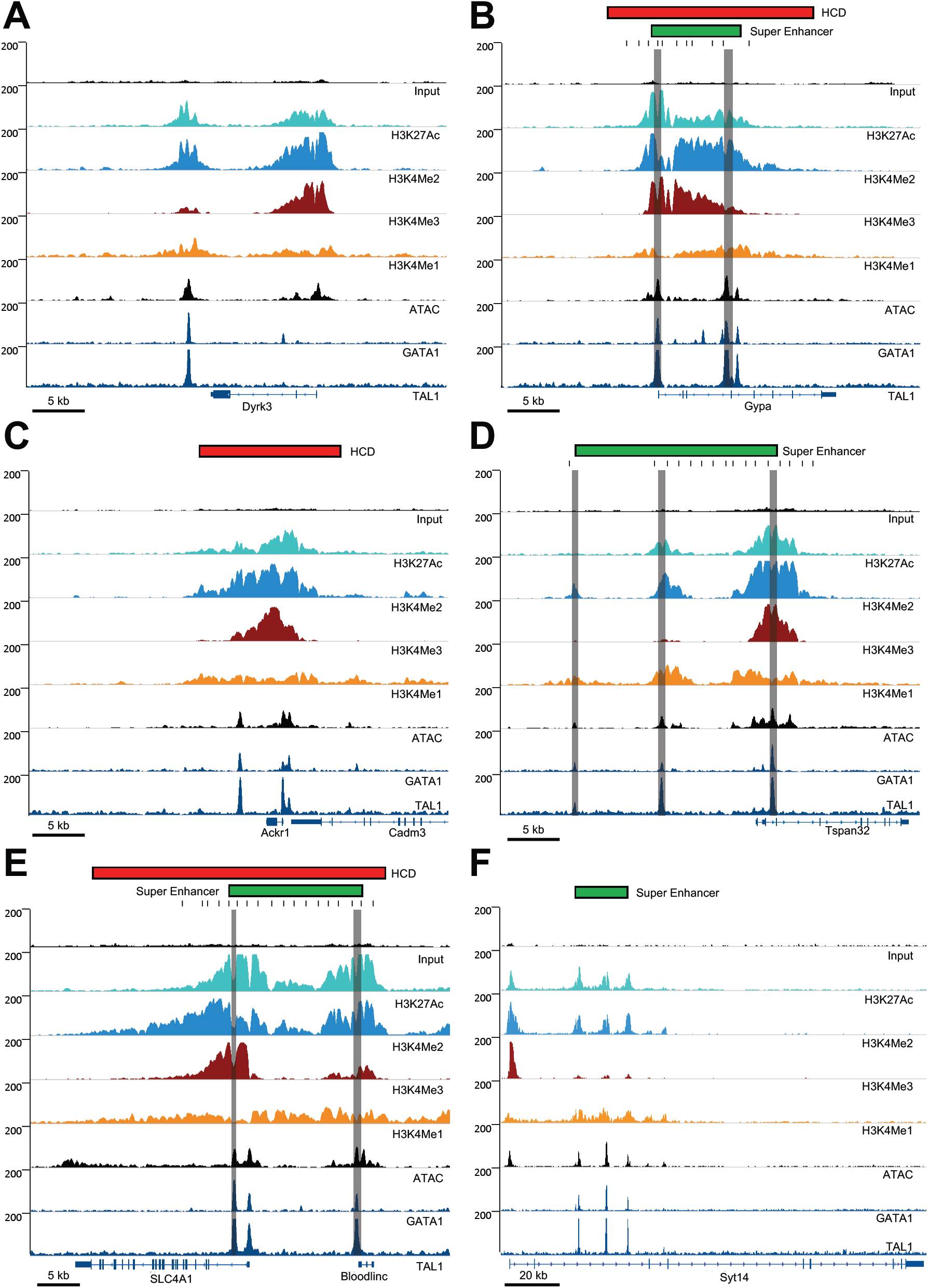
ChIP-seq profiles at selected gene loci. Tracks show read densities for the indicated histone modifications or transcription factors, with additional tracks for the Input control at top and for ATAC-seq. Genes and scale are shown at the bottom, and peak calls for super-enhancers and/or hyperacetylated chromatin domains (HCDs) at the top. Gray shading indicates regions tested by deletion in MEL cells by CRISPR/Cas9-mediated genome editing. Black bars indicate regions amplified in ChIP-qPCR experiments. **(A)** The Dyrk3 locus, harboring a putative enhancer that is called neither a super-enhancer nor an HCD. **(B)** The Gypa locus, harboring both a super-enhancer and an HCD. **(C)** The Ackr1/Cadm3 locus, harboring an HCD but not a super-enhancer. **(D)** The Tspan32 locus, harboring a super-enhancer but not an HCD. **(E)** The SLC4A1/Bloodlinc locus, harboring an HCD and a super-enhancer. **(F)** The Syt14 locus, harboring a super-enhancer but not an HCD.

Our previous definition of hyperacetylated domains involved broad regions of continuous enrichment for specific histone modifications – most often, histone hyperacetylation and H3K4Me2. We therefore analyzed our H3K27Ac and H3K4Me2 ChIP-seq datasets derived from e14.5 murine fetal liver by ranking MACS2^15^ peaks by breadth. The establishment of a formal definition for a domain requires the application of a subjective cutoff value; for our purposes, rather than an absolute peak breadth we chose the top 2% of MACS2 peaks ranked by breadth. We did this because we found that a cutoff based on ranking, as opposed to absolute peak breadth, translated more consistently between different datasets. Our other criterion was that this produced a list of HCDs that was comparable in size to the list of super-enhancers called from the same datasets (see below), and thus facilitated a comparison of the two methods.

To arrive at this list of HCDs, we first intersected the replicates of our ChIP-seqs for each histone modification (H3K27Ac or H3K4Me2), and then applied the 2% cutoff to each intersection, resulting in 420 H3K27Ac and 760 H3K4Me2 peaks. We then identified the H3K27Ac and H3K4Me2 peaks that overlapped within the genome and merged them by taking their union. This resulted in a final tally of 216 regions we term “hyperacetylated chromatin domains” (HCDs) in primary murine fetal liver (Supplementary Fig. 1a, Supplementary File 1).

When choosing criteria to call super-enhancers, we wanted to stay as true to the original method as possible, while also making sure the datasets were comparable to our hyperacetylated domains. We therefore used lineage specific transcription factor (GATA-1 and SCL/Tal1) peaks to identify a set of enhancers in fetal liver^14^. We then ranked these enhancers based on signal intensity in our H3K27Ac and H3K4Me2 ChIP-seqs, since these are the marks that we used to identify hyperacetylated domains. We used the ROSE algorithm^6,7^ with the default stitching distance (12.5 kb) and a +/−500bp TSS exclusion zone, which resulted in 307 super-enhancers ranked on H3K27Ac and 214 super-enhancers ranked on H3K4Me2. To identify super-enhancers for both H3K27Ac and H3K4Me2 we took the union of the 307 and 214 super-enhancers, only where these regions overlapped, for a final tally of 173 murine erythroid super-enhancers. (Supplementary Fig. 1b, Supplementary File 1). Examples of gene loci that exhibit HCDs, super-enhancers, both features, or neither are shown in Figure 1.

Insofar as signal strength has been taken as a proxy for enhancer strength, the fundamental utility of super-enhancer identification has been the ability to pick out associated genes that are presumably the most important factors in cell-type specificity. We therefore compared the properties of genes associated with HCDs to those associated with super-enhancers. For association of genes with super-enhancers, the most common method in the literature is “nearest neighbor” (i.e. nearest gene), and so we used this method to associate super-enhancers with genes. We used the same approximation for HCDs, although notably for 205 out of the 216 regions we have classified as HCDs in murine fetal liver, the nearest active genes are actually located within them. The remaining 11 HCDs all harbor peaks of H3K4Me3, suggesting the presence of unannotated gene promoters (not shown), but for the sake of consistency in this analysis we still associated these HCDs with the nearest annotated genes. Using a published database of Affymetrix-derived gene expression in murine fetal liver (ErythronDB^16,17^), we then compared expression levels for genes associated with HCDs and genes associated with super-enhancers (Figure 2A). We find that HCD-associated genes are expressed at significantly higher levels in murine fetal liver than those associated with super-enhancers.

**Figure 2:**
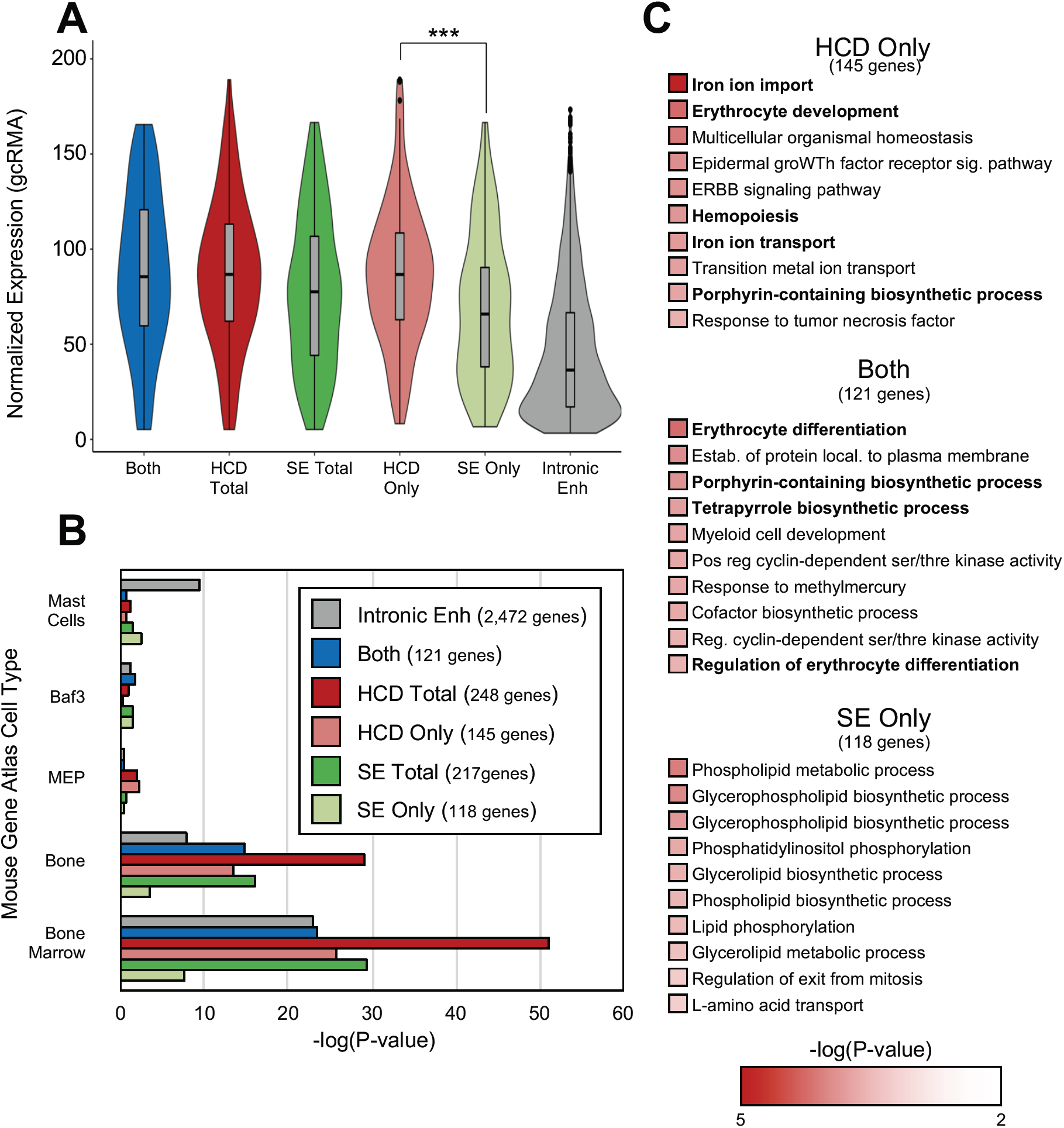
Comparison of genes associated with super-enhancers vs. HCDs in murine erythroid cells. **(A)** Violin plots of expression of genes associated with HCDs, super-enhancers or both. P-values were calculated with a two-tailed Student’s t-test, P=0.00056. Expression for genes associated with all putative enhancers located within introns is shown for comparison. **(B)** Bar graph showing combined score for Enrichr cell-type enrichment for the 5 cell types with the highest scores for each category. **(C)** Listings of the top ten GO terms for biological processes for the indicated groups; erythroid-specific terms are in boldface type.

Not only did we find that HCDs were associated with more highly expressed genes than super-enhancers, but we also found that they were associated with genes that are better able to identify erythroid cell types in the Mouse Gene Atlas than genes associated with super-enhancers, using Enrichr cell type analysis software^18,19^ (Figure 2B, Supplementary File 2). Functional enrichment analysis, also performed through Enrichr, shows that genes associated with HCDs are more likely to identify terms that are specific for erythroid cells. Seven of the top ten most enriched terms for genes associated solely with HCDs are specific for erythroid biology, while in contrast, only two out of the top ten are erythroid-specific using genes associated solely with super-enhancers (Figure 2C, Supplementary File 3). Notably, genes found to be associated with both an HCD and a super-enhancer exhibit a distribution of expression levels similar to those associated with an HCD alone, and thus also higher than those associated only with super-enhancers. In Enrichr analysis, this population of genes also results in the identification of terms more obviously applicable to erythroid biology than with genes associated solely with super-enhancers.

To determine how generalizable our comparison of HCDs and super-enhancers is, we applied the same analyses to a selection of publicly available datasets. We were able to perform domain analysis and super-enhancer analysis on ChIP-seqs derived from human primary cultured CD34+ erythroid cells^20^ in the same fashion as for our murine fetal liver-derived datasets (Figure 3, Supplementary File 4). This analysis resulted in 302 HCDs and 326 super-enhancers, and the same general trend that we see in mouse erythroid cells holds true in human erythroid cells: genes associated with HCDs are both more highly expressed than genes associated with super-enhancers, (Figure 3A, Supplementary File 5) and appear to be more erythroid-specific (Figures 3B,C, Supplementary File 6).

**Figure 3:**
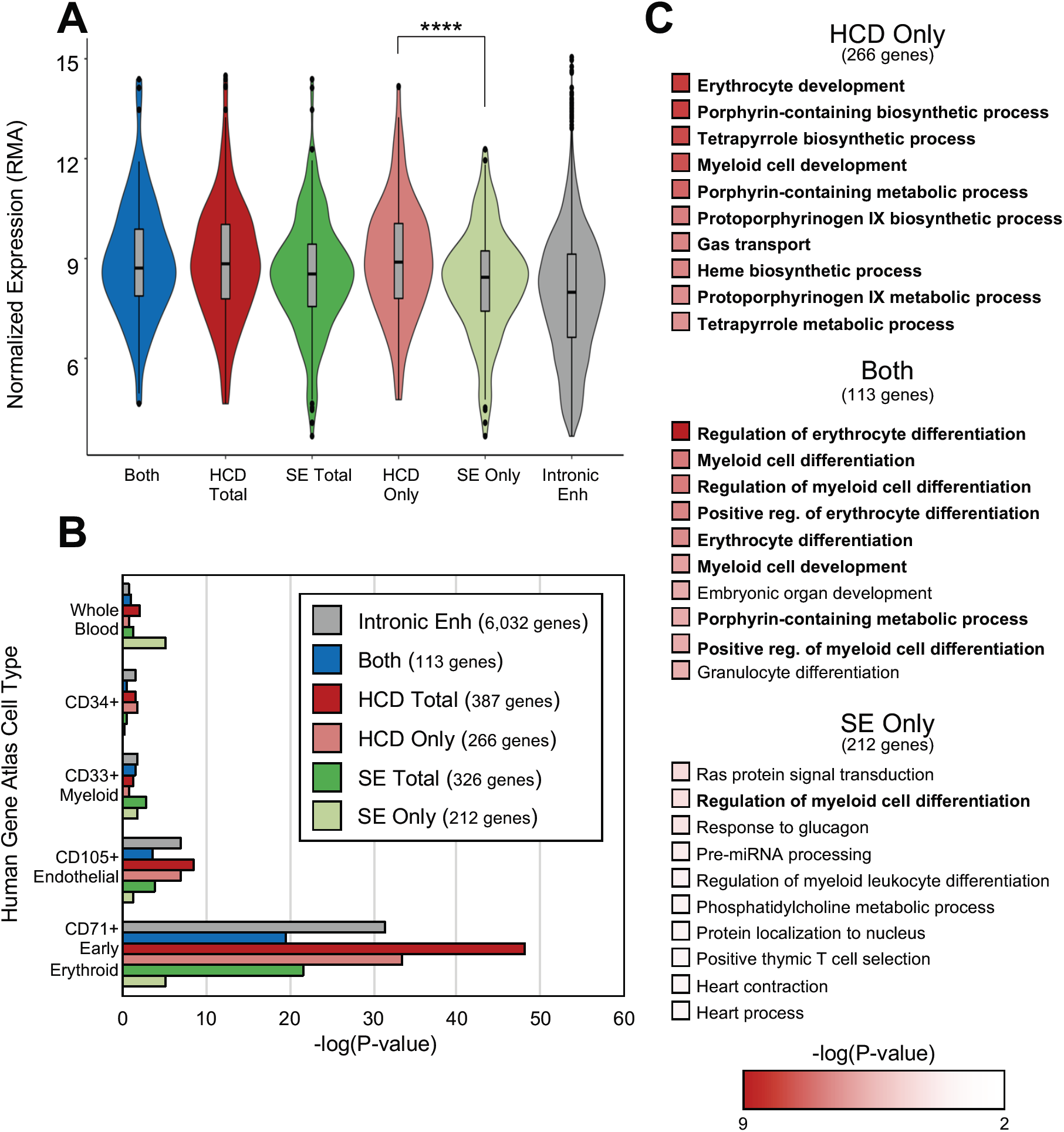
Comparison of genes associated with super-enhancers vs. HCDs in human erythroid cells. **(A)** Violin plots of expression of genes associated with HCDs, super-enhancer or both. P-values were calculated with a two-tailed Student’s t-test, P=0.00073. Expression for genes associated with all putative enhancers located within introns is shown for comparison. **(B)** Bar graph showing combined score for Enrichr cell-type enrichment for the 5 cell types with the highest scores for each category. **(C)** Listings of the top ten GO terms for biological processes for the indicated groups; erythroid-specific terms are in boldface type.

We performed additional comparisons of HCD and super-enhancer analyses using available datasets derived from non-erythroid murine tissues, specifically intestinal epithelium^21^ (Supplementary Figure 3, Supplementary File 7) and retina^22,23^ (Supplementary Figure 4, Supplementary File 8). In these cases, while all rankings are still performed using H3K27Ac and H3K4Me2, enhancer calls for super-enhancer analysis are derived from ATAC-seq instead of TFs, and intestinal epithelial domains were reduced from the top 2% to the top 1% of H3K27Ac/H3K4Me2 peaks ranked by breadth, again to arrive at a list of HCDs comparable in size to the list of super-enhancers (see Materials and Methods). In both cases, the HCDs identify more highly expressed genes (Supplementary Figures 3A, 4A), and genes more closely associated with the specific tissue type, than do super-enhancers (Supplementary Figures 3B-C, 4B-C, Supplementary Files 9-12).

Stratification of ChIP-seq-derived H3K4Me3 peaks by breadth has similarly been demonstrated as a useful tool for the identification of highly-expressed genes important in determining cell identity^24^. We therefore performed a comparison of genes associated with HCDs to genes associated with the broadest 5% of H3K4Me3 peaks in our murine erythroid ChIP-seqs (Supplementary Figure 5). In this comparison, HCDs identified a population of genes that was more highly expressed than that associated with the broadest H3K4Me3 peaks, but the difference was not as great as between HCDs and SEs. Enrichr analysis, however, indicates that H3K4Me3 peaks identify a population of genes more closely associated with erythroid cell types in the Mouse Gene Atlas, along with closer association with erythroid-specific terms in functional enrichment analysis. Notably, however, the set of genes that map within the broadest 5% of H3K4Me3 peaks was considerably larger than that associated with HCDs, and fully 94% of HCD-associated genes fell within this population as well. Thus, this analysis represents a comparison between the set of HCD-associated genes and a larger set that subsumes HCDs.

Our previous characterization of HS-E1 within the β-globin locus as an enhancer required for the formation of an HCD implied that classification of enhancers according to peak breadth could serve to distinguish between enhancers with different functions^12^. To investigate this, we performed CRISPR/Cas9-mediated genetic manipulation of selected gene loci in murine erythroleukemia (MEL) cells^25,26^. These cells are transformed, but exhibit a phenotype similar to proerythroblasts, and they can be induced to mature (but not to enucleate) by incubation in 2% DMSO. Maturation is associated with cessation of cell division in a majority (>90%) of cells and dramatic upregulation of erythroid-specific genes, most notably the β- and β-globin genes.

As an example locus exhibiting an HCD, we chose the region harboring the gene encoding glycophorin A (Gypa), a late-stage erythroid cell surface marker that is highly upregulated upon MEL cell maturation. The region exhibits a substantial domain that encompasses the gene promoter and a pair of putative enhancers located within the third and fourth introns of the gene (Figure 1B). These enhancers, together with the region near (but not directly over) the gene promoter, are also called as a super-enhancer by our ROSE-based analysis. Using a CRISPR/Cas9-based strategy we created a deletion of a 700 bp region encompassing the major GATA1 binding site within intron 3, while leaving exon 4 and the splice acceptor intact.

We measured expression of Gypa in differentiating (DMSO-treated) MEL cells, which more closely resemble e14.5 fetal liver than undifferentiated MEL cells, and saw that Gypa expression decreased >10-fold in cell lines harboring the enhancer deletion (Figure 4A). ChIP-qPCR analysis of the Gypa locus in these cells indicates that deletion of the enhancer has a significant effect on levels of H3K4Me2 (Figure 4B) and H3K27Ac (Figure 4C) observed across the entire region, consistent with a requirement for the enhancer in HCD formation.

**Figure 4:**
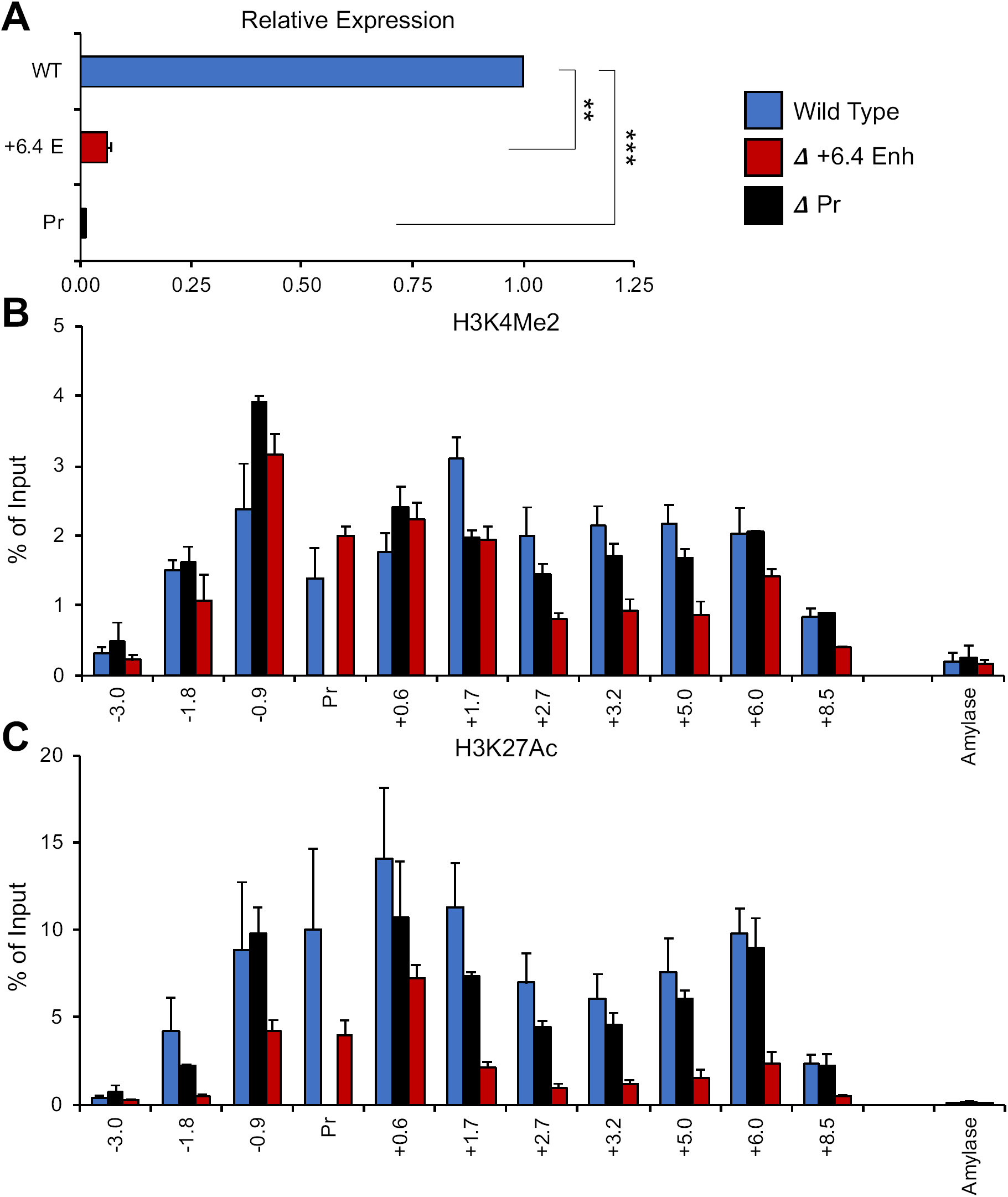
Deletions of the gene promoter and of a putative enhancer within the murine Gypa locus in differentiating MEL cells. **(A)** Bar graph showing the results of quantitative real-time PCR analysis of Gypa expression in wt MEL cells (MEL), cells harboring a deletion of the putative intronic enhancer region (6.4 Enh) or the Gypa gene promoter (Promoter).). P-values were calculated using a two-tailed Student’s t-test;** = P<0.005, *** = P<0.0005. **(B-C)** Bar graph showing percent of input control obtained using PCR probes at the indicated locations (in kb) relative to the transcription start site for the Gypa gene in ChIP assays using antibodies specific for H3K4Me2 **(B)** or H3K27Ac **(C)**. “Amylase” indicates a control probe within the inactive amylase gene locus.

Given that the Gypa HCD occurs largely within the transcribed region of the gene, however, it remains possible that the effect of enhancer deletion on histone modifications is a secondary consequence of decreased transcription. To address this, we created MEL cells harboring a deletion of the Gypa gene promoter. These cells exhibit little to no expression of Gypa (Figure 4A), but levels of histone modifications across the gene are not affected to nearly the same degree as with the enhancer deletion (Figure 4B-C). The results suggest that the histone modifications we have identified as the Gypa HCD are, for the most part, a direct consequence of enhancer activity.

Another region that exhibits an HCD in primary murine erythroid cells is the SLC4A1 locus, harboring the gene that encodes Band 3, an important anion transporter in erythroid cell membranes. The HCD encompasses a large portion of the transcribed region along with sequences extending ∼12 kb upstream of the gene promoter, including a previously defined lncRNA gene, Bloodlinc, that has been reported to be transcribed in erythroid cells^27^. While we detected expression of this lncRNA in the murine fetal liver samples used for our ChIP-seq analyses, we have been unable to detect any expression in MEL cells (not shown). We can identify putative enhancers, as indicated by transcription factor binding, at locations +1.1 kb and −10.4 kb from the SLC4A1 transcription start site; the −10.4 kb enhancer is located near the 3’ end of the annotated Bloodlinc gene.

We evaluated the effects of separate CRISPR/Cas9-mediated deletions of the two putative enhancers within this locus. SLC4A1 is not expressed at measurable levels in proliferating MEL cells, so we confined our analyses to differentiating MEL. Interestingly, individual deletions of either enhancer result in reductions in SLC4A1 mRNA levels (Supplementary Figure 6A), indicating that both enhancers are necessary for normal gene expression. Moreover, both deletions result in substantial decreases in enrichments for H3K27Ac and H3K4Me2 locus-wide (Supplementary Figure 6B+C), with two exceptions: (1) H3K4Me2 at the probe immediately proximal to the SLC4A1 promoter is minimally affected; (2) deletion of the +1.1 kb enhancer has no significant effect on these histone modifications in the immediate vicinity (+/−1 kb) of the −10.4 enhancer. Thus, at this locus, the HCD requires the presence of two enhancers, neither of which is sufficient for domain formation in the absence of the other. As with the Gypa locus, however, the HCD appears to be a consequence of enhancer activity, and is associated with high-level gene expression.

As an example locus exhibiting a super-enhancer, but not an HCD, we investigated the region harboring the gene encoding Tspan32, a member of the tetraspanin superfamily. The super-enhancer identified at this locus includes three putative individual elements, marked by GATA1 binding, located at −17 kb, −10 kb and +0.7 kb relative to the transcription start site (TSS). We created deletions of each of these enhancers individually by CRISPR/Cas9-mediated genome editing in MEL cells. Notably, measurement of Tspan32 gene expression in differentiating MEL cells indicated that only deletion of the +0.7 enhancer had a significant effect (Figure 5A), while deletions of the other two enhancer regions showed no requirement for either element in maintenance of normal steady-state Tspan32 mRNA levels. Moreover, none of the enhancer deletions, including the +0.7 enhancer, showed any effect on histone modifications (H3K27Ac or H3K4Me2) in the region encompassing the Tspan32 gene promoter (Figure 5B-C).

**Figure 5:**
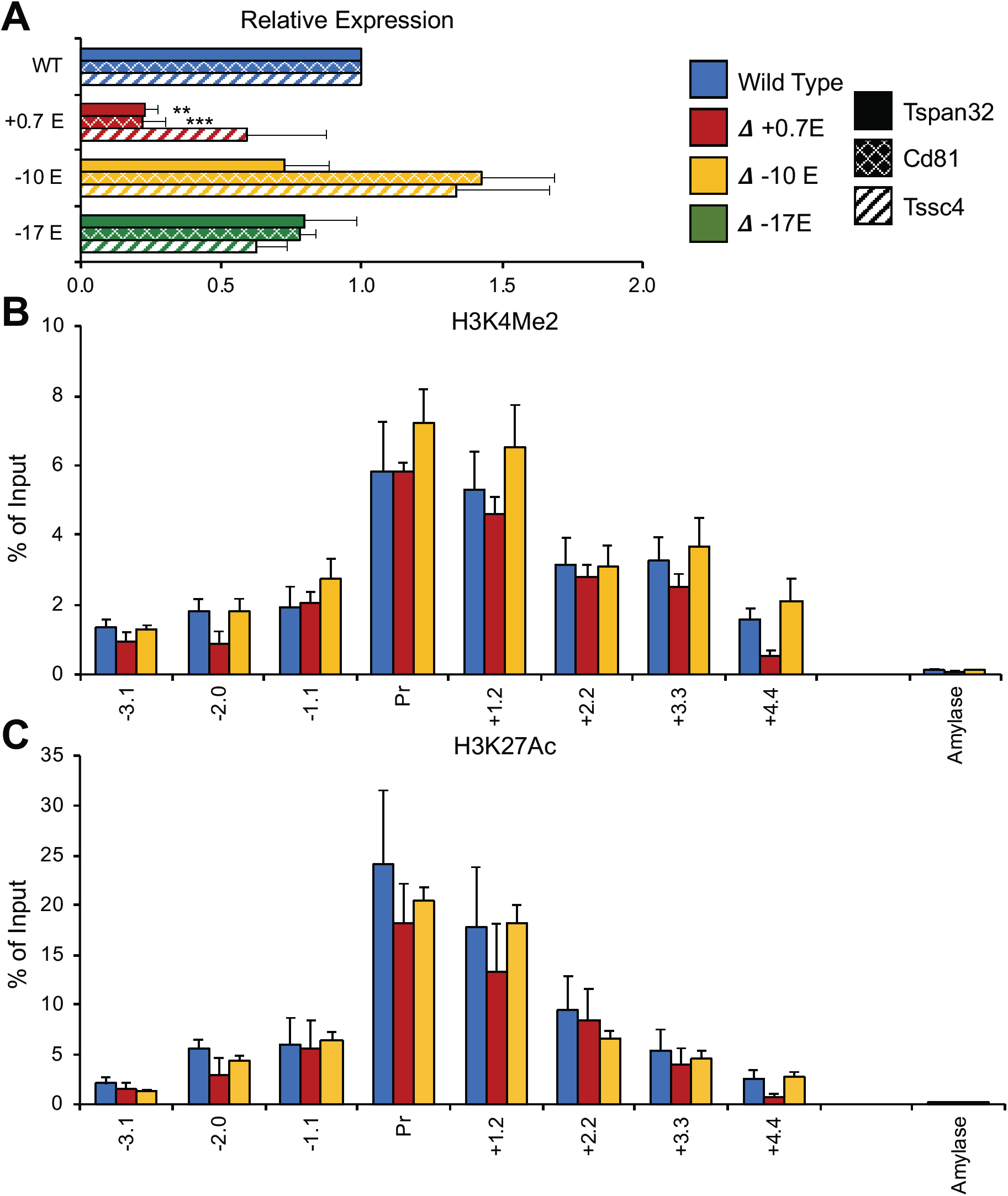
Effects of deletions of putative enhancers in the Tspan32 gene locus in differentiating MEL cells. **(A)** Normalized gene expression for Tspan32, Cd81 and Tssc4 measured by qrt-PCR in wt (MEL) and upon deletion of the indicated enhancer regions (0.7, −10, −17). P-values were calculated using a two-tailed Student’s ttest; * = P<0.05, ** = P<0.005, *** = P<0.0005. **(B-C)** Bar graphs showing % of input control for the indicated probes derived from qrt-PCR analysis of ChIPs using antibodies specific for H3K4Me2 **(B)** or H3K27Ac **(C)**. “Amylase” indicates a control probe within the inactive amylase gene locus.

To further investigate this behavior, we considered the possibility that the “nearest neighbor” assumption – e.g., that the super-enhancer or individual elements within it necessarily regulates only the Tspan32 gene – might not be valid, and so we expanded our analysis to include additional genes neighboring Tspan32. Hi-C data from murine cells^28^ indicate that Tspan32 resides within a topologically associating domain (TAD) that contains two other genes that are expressed in erythroid cells, Cd81 (encoding another tetraspanin superfamily member) and Tssc4 (encoding a protein of unknown function), located 48 and 64 kb from the Tspan32 gene promoter, respectively. Surprisingly, deletion of the +0.7 enhancer had significant effects on the expression of both genes (Figure 5A), which generally paralleled the effects we observed on the Tspan32 gene. Thus, the +0.7 enhancer is required for normal expression of multiple neighboring genes in differentiating MEL cells.

We then utilized a publicly available transcriptomic database (ErythonDB^16,17^) to examine the expression of Tspan32, Cd81 and Tssc4 during normal erythropoiesis. This indicated that Tspan32 is substantially upregulated later in erythroid maturation, while Cd81 and Tssc4 are downregulated. We therefore examined gene expression in our various MEL cell lines prior to differentiation, as an approximation of an earlier stage of erythroid maturation than that modeled by DMSO-treated MEL cells. Interestingly, in this environment, normal expression levels of all three genes required the −10 kb element, while the other enhancers had smaller but still significant effects (Figure 6). This suggests that the components of the Tspan32 super-enhancer have differential activities at different stages of erythropoiesis, with a switch to dependence on the +0.7 kb element from the −10 kb element as maturation proceeds. Notably, ChIP-qPCR analysis of the promoter region of Tspan32 indicates that, as in differentiating MEL cells, none of the enhancer deletions affects histone modifications in undifferentiated MEL cells (Figure 6). This behavior suggests that the specific modifications we have measured over the Tspan32 promoter region are not sufficient for high-level gene expression, and therefore that the Tspan32 enhancers, at distinct maturational stages, function by a different mechanism that does not involve long-range modification of chromatin.

**Figure 6:**
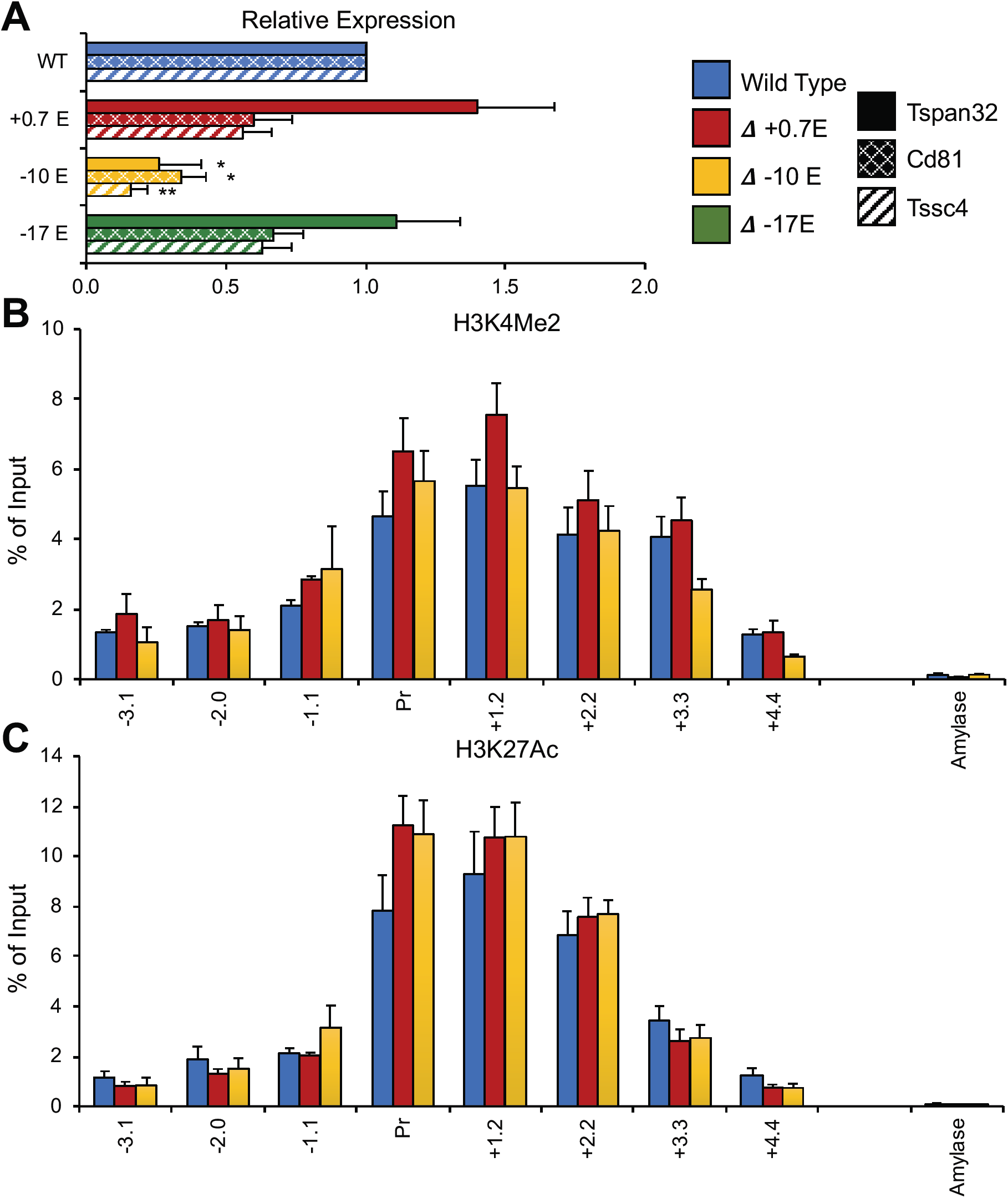
Effects of deletions of putative enhancers in the Tspan32 gene locus in proliferating MEL cells. **(A)** Normalized gene expression for Tspan32, Cd81 and Tssc4 measured by qrt-PCR in wt (MEL) and upon deletion of the indicated enhancer regions (0.7, −10, −17). P-values were calculated using a two-tailed Student’s ttest; * = P<0.05, ** = P<0.005, *** = P<0.0005. **(B-C)** Bar graphs showing % of input control for the indicated probes derived from qrt-PCR analysis of ChIPs using antibodies specific for H3K4Me2 **(B)** or H3K27Ac **(C)**. “Amylase” indicates a control probe within the inactive amylase gene locus.

Genetic manipulations in MEL cells indicate that HCD formation at the Gypa and SLC4A1 gene loci requires the activity of specific enhancers located within them. We therefore asked if the Gypa enhancer was also sufficient to form an HCD. To do this, we employed a “CRISPR-in” strategy, in which a portion of the Gypa enhancer was inserted into the location of the −10 kb element at the Tspan32 locus. This was accomplished using a repair template consisting of the Gypa enhancer sequence flanked by homology arms, targeted to the site of the deleted Tspan32 −10 kb element by specific gRNAs. We find that in subclones homozygous for this insertion, expression of the Tspan32 gene increases in differentiating MEL cells (Figure 7), where the Gypa enhancer is most active. Notably, the +0.7 kb element, which is also active in differentiating MEL, is still present in these cells, and so this represents a super-activation over normal transcription levels in this cellular context.

**Figure 7:**
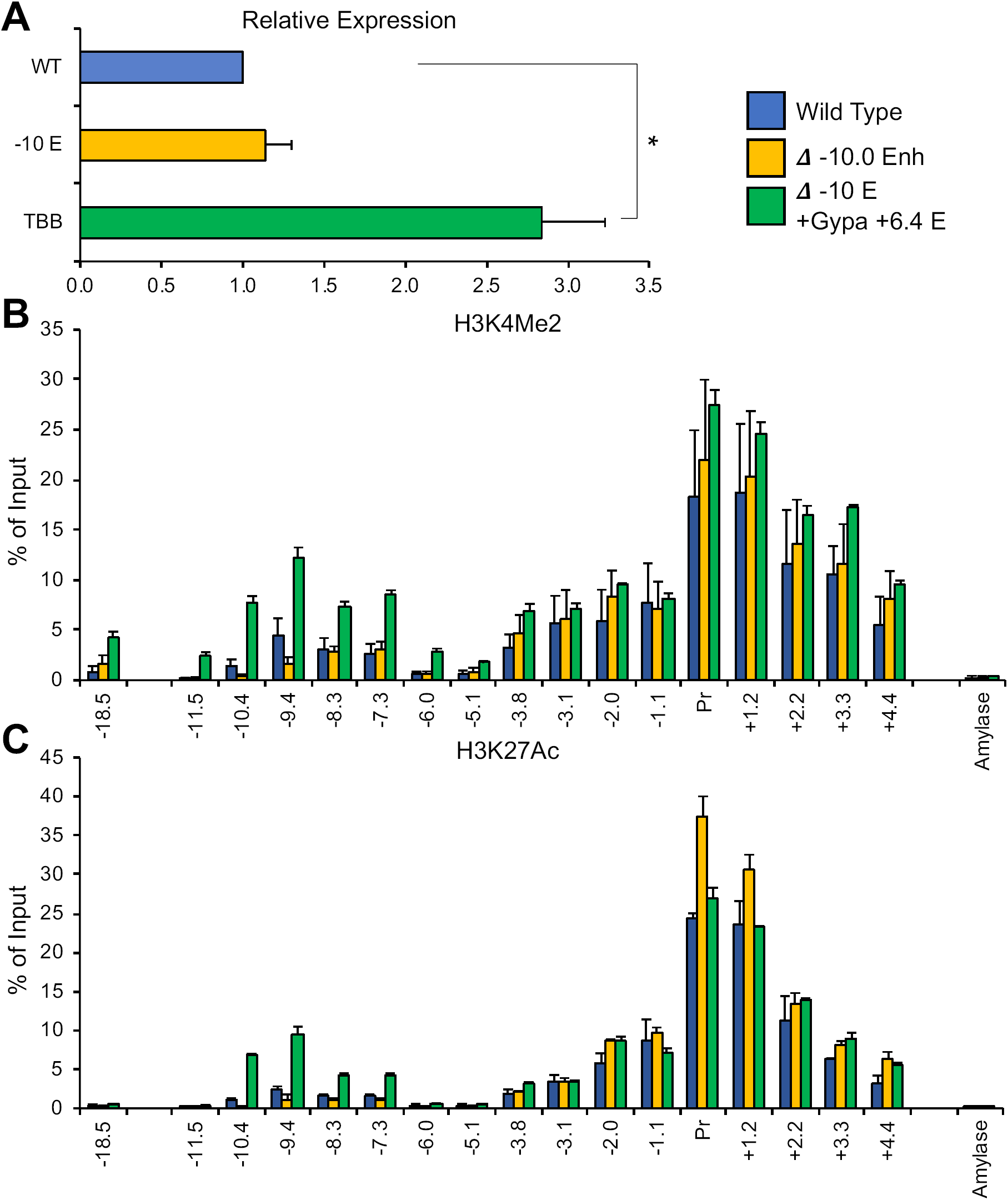
Effects of addition of the Gypa +6.4 enhancer in the Tspan32 gene locus in differentiating MEL cells. **(A)** Normalized gene expression for Tspan32 measured by qrt-PCR in wt (MEL) and upon deletion of the - 10.0 enhancer or insertion of the Gypa +6.4 enhancer. P-values were calculated using a two-tailed Student’s ttest; * = P<0.05. (B+C) Bar graphs showing % of input control for the indicated probes derived from qrt-PCR analysis of ChIPs using antibodies specific for H3K4Me2 **(B)** or H3K27Ac (C). “Amylase” indicates a control probe within the inactive amylase gene locus.

We also find, however, that introduction of the Gypa enhancer results in significant increases in H3K27Ac and H3K4Me2 at the site of insertion and for substantial distances in either direction, including probes located between 5-10 kb upstream. Thus, the Gypa enhancer is sufficient to induce the formation of an HCD when introduced into the Tspan32 locus.

## Discussion

Ranking of enhancers by “strength” as determined by ChIP-seq signal intensity is now a highly common technique for meta-analysis of epigenomic data, with applications in revealing cell type-specific pathways and transcriptional regulatory networks. In this regard, the super-enhancer model has generated a great deal of interest, especially insofar as it can be applied to dysfunctional or disease states. In this study, however, we have demonstrated that for the specific purposes of identifying the genes and pathways most important for cell phenotype, a ranking according to peak breadth of regions identified by ChIP-seq using antibodies for two commonly assayed histone modifications is more useful. The combination of H3K4Me2 and H3K27Ac “domains” – both of which are detected using widely available and highly robust antibodies – serves to identify genes that are both more highly expressed and more closely aligned to cell type than genes identified by super-enhancer analysis. Moreover, the protocol for classifying regions by peak breadth and identifying HCDs, which we include here (Supplemental Figure 1, Supplementary Files 13-14) is both more transparent and simpler to use than the ROSE algorithm.

Notably, peak breadth in histone modification ChIP-seqs has previously been used to identify highly-expressed and/or cell type-specific genes. “Stretch enhancers,” for example, have been defined as regions exhibiting continuous epigenomic signatures (“chromatin state” from ChromHMM^29^) indicative of enhancers over spans of 3 kb or more^30^. This contrasts with our own analysis in two key respects. First, hyperacetylated domains encompass both enhancers and promoters and are not necessarily limited to either type of sequence element. Second, the “stretch enhancer” definition applies to fully 10% of all putative enhancer regions, while our own definition of a hyperacetylated domain is more restrictive.

There are several potential explanations for the differences in genes and pathways identified by HCD (peak breadth) vs. super-enhancer analysis. Assignment of genes to super-enhancers relies on the assumption that a given enhancer regulates the nearest active gene. This is, at best, an approximation, the accuracy of which is impossible to evaluate in the absence of comprehensive genetic analysis or other data, and difficult even then. In contrast, genes assigned to HCDs are nearly always (95%) located within them, and there is a logically greater certainty that a gene within an HCD is directly affected. Complexity arises in those cases involving multiple active genes within an HCD, and so even the domain concept involves a degree of approximation, but based on gene expression and cell type-specificity (Figures 2-3), the uncertainty involved appears to be less than that inherent to super-enhancers.

To a certain extent, however, HCDs may be advantaged compared to super-enhancers due to purely technical aspects of ChIP-seq analysis. Ranking by peak intensity (e.g. peak height) is limited because mapping algorithms routinely discard duplicate reads as a precaution against PCR artifacts arising from library construction. Reads therefore cannot stack on top of each other, and the maximum height a given peak can achieve is artificially limited. Ranking by peak breadth avoids this necessary limitation and therefore may present a more sensitive metric for determining the strongest and/or most cell type-specific enhancers.

An important consideration is the potential mechanisms that underlie differences in peak intensity vs. differences in peak breadth. In ChIP-seqs, which assay large numbers of cells at once, differences in peak intensity most likely reflect differences in binding affinity between enhancer regions; thus, a higher peak intensity for a transcription factor reflects a higher proportion of alleles crosslinked to that factor (whether directly or indirectly), and for histone modifications a higher proportion of alleles associated with the cognate enzyme(s). In either case, binding does not necessarily invoke transcriptional activation of a neighboring gene.

In contrast, the largest peak breadths most likely arise from a mechanism distinct from higher binding affinity. In our previous investigation of the HCD within the murine β-globin locus, we demonstrated that the association of specific histone modifications was not intrinsic to the entire sequence underlying the HCD, but was a function of a smaller enhancer region within it. We demonstrate similar effects for enhancers within the Gypa and SLC4A1 gene loci in this study. This requirement strongly suggests that the histone modification pattern associated with an HCD spreads in some fashion from the regulatory element(s). Regardless of the mechanism that underlies such spreading – whether a specific mechanism akin to the formation of heterochromatic domains, or simple “spill-over” from an excess of chromatin modifiers – a broad domain implies a high degree of binding and activity, and insofar as the vast majority of HCDs subsume gene promoters, of activity at transcription initiation sites. Rather than implying that a greater proportion of alleles in the cell population harbor modified nucleosomes, an HCD implies a greater absolute quantity of modified histones at each allele, which could conceivably contribute to stability of the active transcriptional state in the nuclear environment.

Notably, ranking of H3K27Ac/H3K4Me2 peaks by breadth is potentially useful in identifying enhancers that function by different mechanisms. In this study, we contrast the effects of deletion of enhancers from within the endogenous Gypa, SLC4A1, and Tspan32 gene loci, which are all classified as super-enhancers, but only the Gypa and SLC4A1 loci as HCDs. At all loci, enhancer deletions result in substantial losses in gene expression levels. At the Gypa and SLC4A1 loci, deletion of the enhancers results in loss of the broad peak of histone hyperacetylation, including substantial losses in modification levels associated with the region proximal to the TSS. At the Tspan32 locus, however, deletion of neither the −10 kb element in undifferentiated MEL cells nor the +0.7 element in differentiating MEL cells results in any effect on chromatin structure at gene promoters or the regions surrounding them. Moreover, introduction of the Gypa enhancer into the Tspan32 locus, at the site of the deleted −10 kb element, results in the formation of an HCD over a region of the locus that normally does not exhibit one. The data suggest a functional distinction between enhancers that work, at least in part, *via* modulation of long-range chromatin structure, and enhancers that do not.

The relationship between enhancer function and HCD formation, however, does not appear to follow the simplest possible model. Both of the putative enhancers within the SLC4A1 locus are required for HCD formation, despite a separation of nearly 12 kb between them. This collaborative function raises the question of whether HCD formation by the Gypa enhancer inserted within the Tspan32 locus is fully enhancer-autonomous, or if instead it requires additional elements that are present both at this locus and at its native location. Additional genetic manipulations will be required to address this question.

An additional concern is the “stitching” step of super-enhancer analysis, in which putative enhancer regions located within a specific distance of each other (the default in the ROSE algorithm is 12.5 kb, and so this is most often used) are considered as a single element, under the assumption that closely-spaced enhancers work together in activation of the same gene or genes^6–8^. Some studies have questioned the general validity of this assumption^10,31^, and our own analysis of the Tspan32 locus presents a clear example of how the super-enhancer concept tends to oversimplify more complex patterns of gene regulation. We found that the Tspan32 super-enhancer, comprised of three distinct candidate enhancer regions, is in fact involved in the regulation of two additional genes as well as Tspan32. Moreover, individual enhancers within the super-enhancer exhibit different roles, with the −10 kb enhancer required solely in undifferentiated MEL cells, the +0.7 kb enhancer required solely in differentiating MEL cells, and the −17 kb enhancer showing no requirement at all. Based on such behavior, the classification of these three elements as a single super-enhancer appears to obscure the functions of these regions more than it illuminates them.

While super-enhancers can identify highly expressed genes that characterize cell identity, we find that HCDs define a set of genes that exhibit higher expression, and better specify cell lineage. Furthermore, as an analytical tool, HCDs present additional advantages over super-enhancers, including a significantly simpler workflow. Our results suggest that super-enhancers may not have a function distinct from that of typical enhancers, and that clustering of enhancer elements does not necessarily imply cooperative function, whereas there appears to be a functional difference between at least some of the enhancers located within HCDs, involving modulation of long-range chromatin structure, compared to other enhancers.

## Acknowledgements

The authors thank Laurie Steiner and Patrick Murphy for critical reading of the manuscript.

## Materials & Methods

### Mice and tissues

Mice (C57BL/6J) were mated overnight and the vaginal plug verified the next morning, indicating embryonic day 0.5 (e0.5). At e14.5 pregnant mice were killed by cervical dislocation for collection of fetal liver. Animal experiments were approved by the University of Rochester Committee on Animal Resources. For ATAC-seq (see below), E14.5 liver-derived proerythroblasts were sorted as previously described^32^, but also including Cd117 staining using a FACS Aria-II. Briefly, larger single cells that were Ter^lo^ CD117+CD44^hi^ were collected. These cells are also positive for CD71.

### ChIP-seq

Chromatin immunoprecipitation was performed as previously described^33^. Briefly, 2 × 10^7^ cells were washed with PBS once, then cross-linked with 1% formaldehyde for 10 minutes at room temperature. Cross-linking was quenched with 5 M glycine for 1 minute at room temperature, and cells were washed with PBS once. Cells were incubated in swelling buffer for 20 minutes on ice, followed by dounce homogenization to isolate cross-linked nuclei. Nuclei were placed in lysis buffer for 30 minutes and then sonicated into ∼200 bp fragments using a Diagenode Bioruptor. Samples were diluted and immunoprecipitated with an antibody to H3K4Me1 (Abcam #ab8895), H3K4Me3 (Active Motif #39916), H3K27Ac (Active Motif #39134), H3K4Me2 (Millipore #07-030), or nonspecific rabbit immunoglobulin G (Millipore #12-370) on a rotator for 18 hrs. at 4°C. DNA-protein complexes were recovered with protein G magnetic beads (Invitrogen). Library preparation was performed as previously described^34^. Library quality was evaluated on a Bioanalyzer and sequencing was performed on a HiSeq 2500 Rapid Run to obtain 1×50bp reads.

### ChIP-Seq Analysis

Illumina reads were converted to the fastq format using bcltofastq-1.8.4 with default parameters. Quality control and adapter removal was performed using Trimmomatic-0.32^35^ using the following parameters “SLIDINGWINDOW:4:20 TRAILING:13 LEADING:13 ILLUMINACLIP:adapters.fasta:2:30:10 MINLEN:15”. All quality reads were aligned to the mm9 reference genome using Bowtie-1.0.1^36^, suppressing multi-mapping reads using the ‘-m 1’ parameters. All alignments were written in the SAM format (-S), converted to BAM and sorted for all subsequent analyses using samtools^37^. Peaks were called for each replicate of each mark using MACS2 along with additional settings including (--broad –broad-cutoff 0.1 −B) using the total input control as the mock data file^16^. The intersection of each replicate was identified using bedtools intersect and was used for subsequent analyses^38^.

### ATAC-Seq

ATAC-seq was performed as previously described^39^. Briefly, 5 × 10^4^ sorted proerythroblasts were lysed by gently pipetting in cold lysis buffer. Cell lysate was resuspended in transposition reaction mix (Illumina) and incubated at 37°C for 30 min. Reactions were purified using AmpureXP beads (Beckman Coulter) following manufacturer’s protocol with minor changes. Beads were used at a 1:1.1 ratio and reactions were washed twice. After purification samples were amplified using 1×NEBnext PCR master mix and 1.25 μM of custom Nextera PCR primers 1 and 2^40^. Libraries were amplified again for an additional 17-19 cycles and left side size selected with SPRIselect beads (Beckman Coulter) at a 1:1 ratio following manufacturer’s protocol, then right side size selected with SPRIselect beads (Beckman Coulter) at a 1:0.5 ratio following manufacturer’s protocol. Library quality was evaluated on a Bioanalyzer and sequencing was performed on a Hiseq2500v2 platform in rapid mode to generate 50 million reads per sample.

### ATAC-seq Analysis

Illumina reads were converted to the fastq format using bcltofastq-1.8.4 with default parameters. Quality control and adapter removal was performed using Trimmomatic-0.32^35^ using the following parameters “SLIDINGWINDOW:4:20 TRAILING:13 LEADING:13 ILLUMINACLIP:adapters.fasta:2:30:10 MINLEN:15”. All quality reads were aligned to the mm9 reference genome using Bowtie-1.0.142^36^, suppressing multi-mapping reads using the ‘-m 1’ parameters. All alignments were written in the SAM format (-S), converted to BAM and sorted for all subsequent analyses using samtools^37^. Reads aligning to organism-specific blacklist regions and the mitochondrial genome are discarded. Accessible regions are identified using MACS2^15^ and ATAC specific parameters (--nomodel --shift −100 --extsize 200).

### RNA-Seq Analysis

Raw fastq files for publicly available datasets (GSE87064-SRR4253101, SRR4253102, GSE98724-SRR5520174 SRR55201745) were downloaded using fastqDump available in SRAtoolkit^21–23^. All reads were processed using trimmomatic^35^ (v0.36) to remove low quality bases and any residual adapter sequence (TRAILING:13 LEADING:13 ILLUMINACLIPtrimmomatic_adapters.fasta:2:30:10 SLIDINGWINDOW:4:20 MINLEN:15). Quality reads were aligned to either mm9/hg19 (depending on organism of origin) using STAR^41^ (v2.5.2b) (STAR --twopassMode Basic --readFilesCommand zcat --runThreadN --runMode alignReads --genomeDir --readFilesIn --outSAMtype BAM Unsorted --outSAMstrandField intronMotif --outFileNamePrefix --outTmpDir --outFilterIntronMotifs RemoveNoncanonical --outReadsUnmapped Fastx). Read alignments were quantified using featureCounts (v1.5.0-p3) (-s 2 -T -t exon -g gene_name -a genes.gtf –o) and normalization was performed within the R (3.4.1) framework using DeSeq2 (v1.16.1)^42^.

### HCD Analysis

The top 2% broadest peaks were filtered from the H3K4Me2 and H3K27Ac intersected replicates. Bedtools intersect was used to identify the intersection of these two tracks, which is what we define as a hyperacetylated domain^36^.Code to call HCDs is included in the supplementary files (identify_HAD.pl). HCDs were associated with nearby genes using bedtools closest, default settings. Microarray expression data for e14.5 fetal liver (maturational stages proerythroblast, basophilic erythroblast, and polychromatic erythroblast) available through ErythronDB (http://www.cbil.upenn.edu/ErythronDB/resources.jsp) were downloaded and used to associate nearby gene expression to domains and non-domains^16,17^. Enrichr (http://amp.pharm.mssm.edu/Enrichr/) was used to perform comprehensive enrichment analysis on associated genes^18,19^.

### Super-Enhancer Analysis

For the identification of super-enhancers within our H3K4Me2 and H3K27Ac datasets we used ROSE (http://younglab.wi.mit.edu/super_enhancer_code.html), created by the Young lab^6^. Input enhancers were defined as a merged peak set of all Gata1 and Tal1 replicates and a TSS_EXCLUSION_ZONE_SIZE of 500bp. Super-enhancers were associated with nearby genes using bedtools closest, default settings. Enrichr (http://amp.pharm.mssm.edu/Enrichr/) was used to perform comprehensive enrichment analysis on associated genes^18,19^.

### Me3 Domain Analysis

The top 5% broadest peaks were filtered from the H3K4Me3 intersected replicates. Domains were associated with nearby genes using bedtools closest, default settings. Enrichr (http://amp.pharm.mssm.edu/Enrichr/) was used to perform comprehensive enrichment analysis on associated genes^18,19^.

### Analysis of Human ChIP-Seq and Microarray Data

Domains and super-enhancers were identified and evaluated during human fetal erythropoiesis using publicly available data (GSE36985)^20^. Mapped read bed files were downloaded and converted to the BAM format (bedtobam bedtools) for fetal H3K4Me2, H3K27Ac, Gata1, Tal1 and the total input control samples. MACS2 was used to identify histone marks (-B –broad –broad-cutoff 0.1) and transcription factor peaks (-B –q 0.01) relative to the total input control^15^. Domains were defined as the intersect of the largest 2% H3K4Me2 (regions larger than 8,015 bp, 936 regions) and H3K27ac (regions larger than 9,020 bp, 503 regions), for a total of 302 regions. Two super-enhancer populations were called using ROSE and identified based on a Gata1/Tal1 union enhancer population which was ranked on H3K4Me2 or H3K27Ac^6^. The resulting sets of super-enhancers were intersected to identify a final population of super-enhancers (326). Domains and super-enhancers were associated with hg18 RefSeq Genes (UCSC) based on a nearest neighbor analysis using bedtools closest (default settings)^33^. Enriched cell types were evaluated using Enrichr and the Human Gene Atlas. RNA expression log2(RMA) were downloaded from ArrayExpression (G_GEOD-36984)^18–20^.

### CRISPR Deletion and Insertion

px458 plasmids harboring sgRNAs targeting specific regions of the mouse genome were engineered as previously described^43^ (see Supplementary Table 1). For deletions MEL745a cells were transfected with one plasmid targeting upstream and one plasmid targeting downstream sequence surrounding the enhancer using Lipofectamine 3000 (Invitrogen) according to manufacturer instructions. For insertions MEL745a cells were transfected with one plasmid targeting the sequence surrounding the previously deleted enhancer and a double stranded repair template using Lipofectamine 3000 (Invitrogen) according to manufacturer instructions. At 48 hrs. the cells were resuspended in D-PBS (Gibco) +0.5% FBS(Gemini), stained with DAPI(ThermoFisher), and the viable, GFP positive population was sorted in bulk. Seven days after sorting, cells were diluted to 1 cell/100μl in Dulbecco modified Eagle medium (Gibco) containing 20% FBS, 1% Glutamax (Gibco), and 1% Pen/strep (Gibco) and plated in a 96 well plate. Homozygous deletions were identified through PCR of the region surrounding the targeted enhancer sequence, and PCR products were Sanger sequenced for confirmation.

### Cell Culture

Cell lines were maintained at 37°C in a CO_2_-humidified atmosphere. MEL745a cells were cultured in Dulbecco modified Eagle medium containing 10% FBS, 1% Glutamax, and 1% Pen/strep. For induction, 2% dimethyl sulfoxide (Fisher) was added to the culture medium.

### ChIP-qPCR

Cells were formaldehyde crosslinked, sonicated and immunoprecipitated, and DNA isolated as for the ChIP-sequencing, with minor differences. Nuclei were isolated from 1 × 10^7^ cells for each experiment and then sonicated to ∼500 bp fragments of genomic DNA using a Diagenode Bioruptor. Samples were diluted and immunoprecipitated with antibodies specific for H3K27Ac (Active Motif # 39134), H3K4Me2 (Millipore # 07-030), and nonspecific rabbit immunogloblulin G (Milipore # 12-370). Analysis was performed using qPCR and detected using the CFX Connect Real-Time PCR System and CFX Manager (Bio-Rad). Primers were designed to amplify regions within the Glycophorin A locus or Tspan32 locus, and as a negative control the Amylase gene promoter (a region that is inactive in erythroid cells)(see Supplementary Table 2).

### qRT-PCR

RNA was isolated using TRIzol (Invitrogen) according to manufacturer instructions. One microgram of RNA was used to synthesize cDNA using the iScript cDNA synthesis kit (Bio-Rad). cDNA was amplified using iTaq Universal SYBR Green Supermix (Bio-Rad) and detected using the CFX Connect Real-Time PCR Sytem and CFX Manager (Bio-Rad). Primers were designed to amplify Gypa, Tspan32, Cd81, and Tssc4 cDNAs, and also ribosomal 18S as a control (see Supplementary Table 3).

### Statistics

Gene expression and ChIP-qPCR experiments were performed on three biological replicates with the exception of the enhancer replacement experiment where only two biological replicates of the homozygous insertion of the new enhancer were found. (standard error (s.e.) is shown. Statistical significance for gene expression was calculated using a two-tailed Student’s t-test.

## Data Availability

All ChIP-seq and ATAC-seq data are available through GEO: GSE132130.

---

**Supplementary Figure 1:**
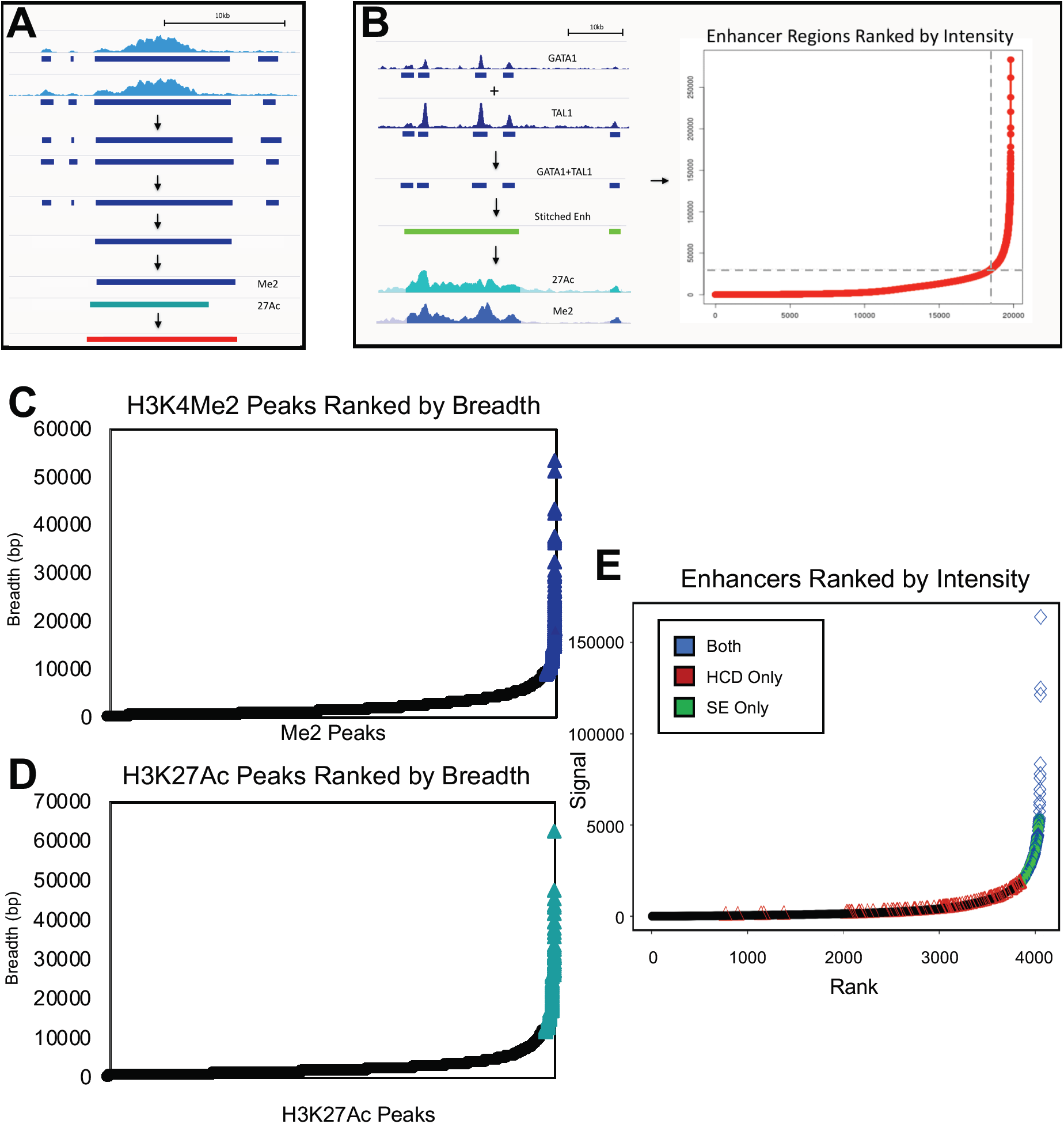
Identification of hyperacetylated chromatin domains (HCDs) and superenhancers. **(A)** Flow chart for HCD calls. MACS2 was used to call peaks in ChIP-seq tracks for H3K4Me2 and H3K27Ac. For each modification, 2 replicate ChIP-seqs were performed, and so the intersection of MACS2-called peaks was taken and then filtered for the top 2% broadest regions. The final list of HCDs was defined as the union of top 2% peaks that overlapped for each modification. **(B)** Flow chart and signal strength ranking graph for super-enhancer calls. A population of putative enhancers was called based on the union of MACS2-called peaks from GATA1 and TAL1 ChIP-seqs. The resulting list was then input into the ROSE algorithm with a stitching distance of 12.5 kb and an exclusion of +/− 500 bp around known TSSs, and ranking based on H3K27Ac reads from our own ChIP-seqs at the selected regions. **(C)** Graph of all H3K4Me2 peaks as ranked by breadth, with K4Me2 domains highlighted in blue. **(D)** Graph of all H3K27Ac peaks as ranked by breadth, with K27Ac domains highlighted in aqua. **(E)** Graph of all H3K27Ac peaks as ranked by ROSE, with super-enhancers highlighted in green, HCDs in red and regions called as both HCDs and super-enhancers in blue.

**Supplementary Figure 2:**
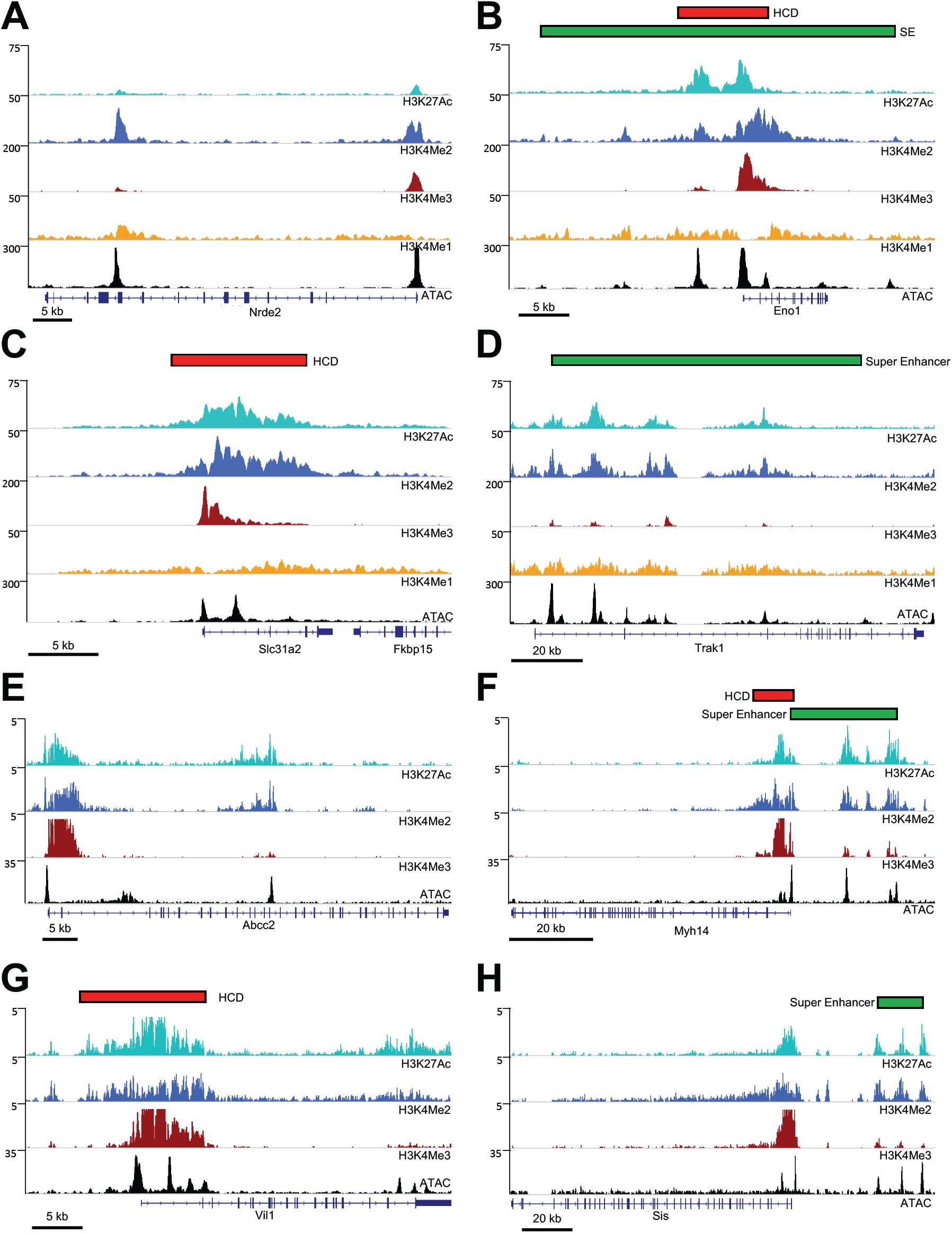
ChIP-seq profiles at selected gene loci in mouse retina (A-D) and intestinal epithelial cells (E-H). Tracks show read densities for the indicated histone modifications or ATAC-seq. Genes and scale are shown at the bottom, and peak calls for super-enhancers and/or hyperacetylated chromatin domains (HCDs) at the top **(A)** The Nrde2 locus, harboring a putative enhancer that is called neither a superenhancer nor an HCD. **(B)** The Eno1 locus, harboring both a super-enhancer and an HCD. **(C)** The Slc31a2/Fkbp25 locus, harboring an HCD but not a super-enhancer. **(D)** The Trak1 locus, harboring a superenhancer but not an HCD. **(E)** The Abcc2 locus, harboring a putative enhancer that is called neither a superenhancer nor an HCD. **(F)** The Myh14 locus, harboring both a super-enhancer and an HCD. **(G)** The Vil1 locus, harboring an HCD but not a super-enhancer. **(H)** The Sis locus, harboring a super-enhancer but not an HCD.

**Supplementary Figure 3:**
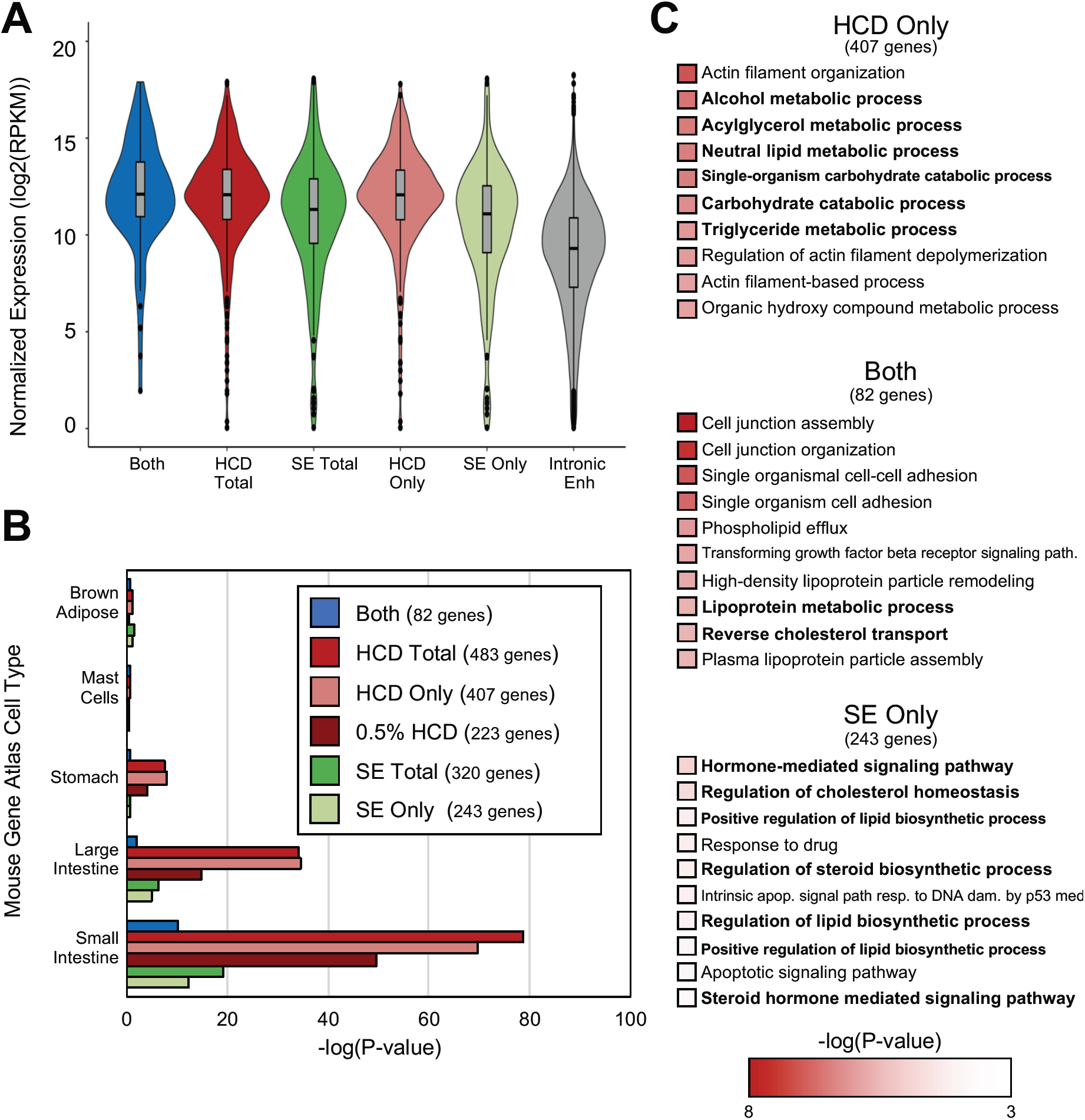
Comparison of genes associated with super-enhancers vs. HCDs in murine intestinal epithelial cells. (A) Violin plots of expression of genes associated with HCDs, super-enhancers or both. Expression for genes associated with all putative enhancers located within introns is shown for comparison. **(B)** Bar graph showing combined score for Enrichr cell-type enrichment for the 5 cell types with the highest scores for each category. **(C)** Listings of the top ten GO terms for biological processes for the indicated groups; intestine-specific terms are in boldface type.

**Supplementary Figure 4:**
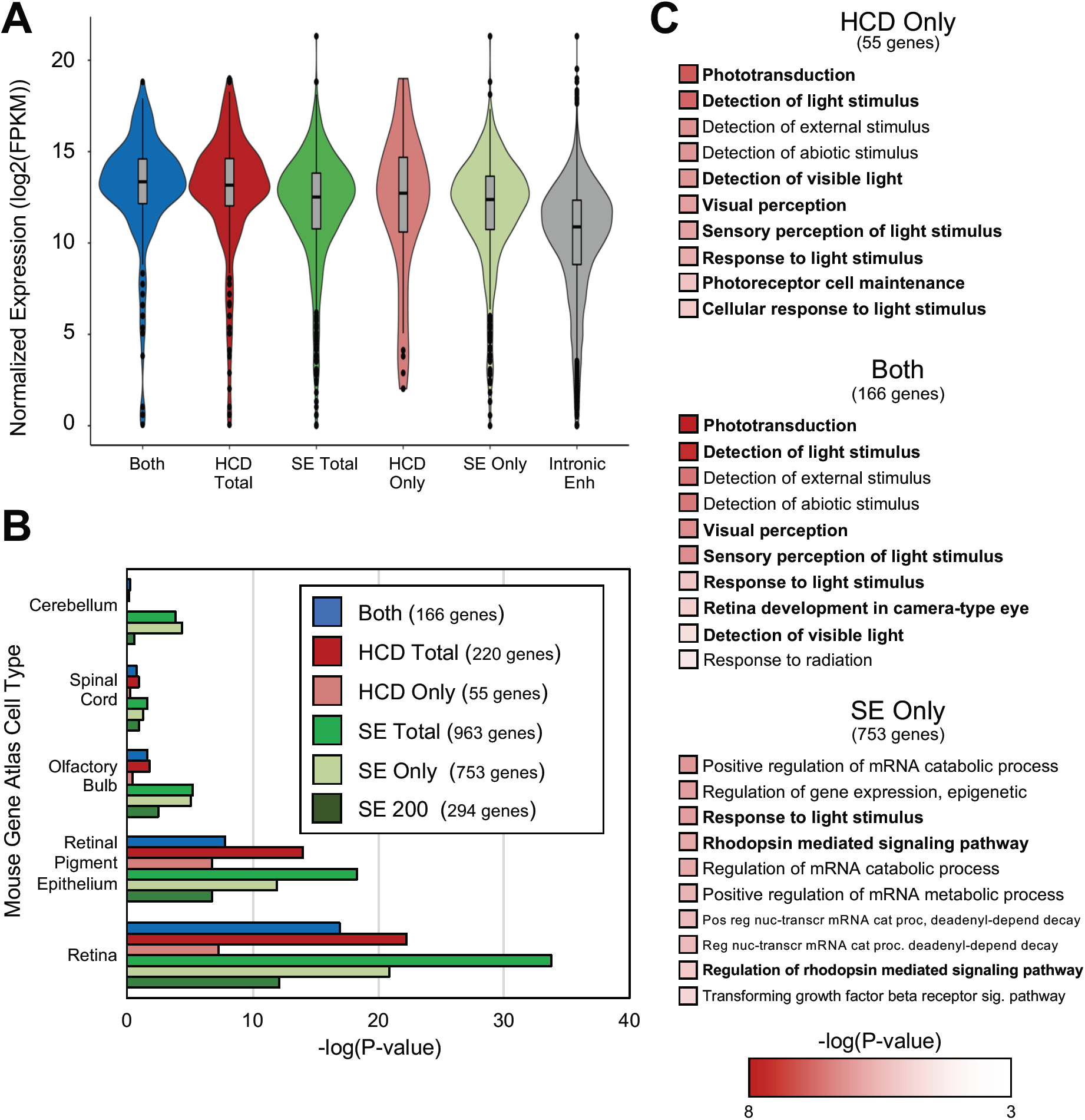
Comparison of genes associated with super-enhancers vs. HCDs in murine retinal cells. **(A)** Violin plots of expression of genes associated with HCDs, super-enhancers or both. Expression for genes associated with all putative enhancers located within introns is shown for comparison. **(B)** Bar graph showing combined score for Enrichr cell-type enrichment for the 5 cell types with the highest scores for each category. **(C)** Listings of the top ten GO terms for biological processes for the indicated groups; retina-specific terms are in boldface type.

**Supplementary Figure 5:**
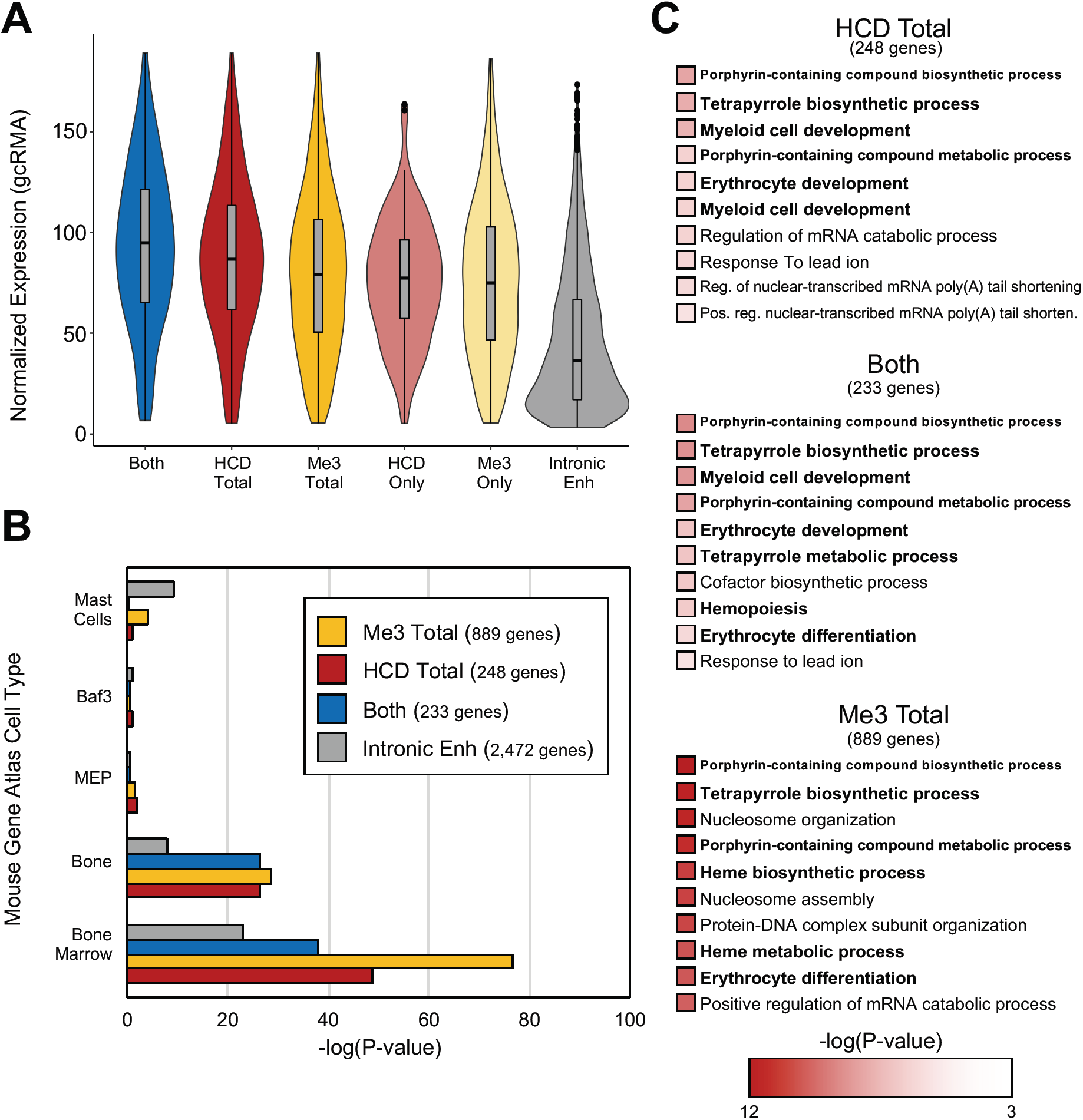
Comparison of genes associated with Me3 domains vs. HCDs in murine erythroid cells. **(A)** Violin plots of expression of genes associated with HCDs, Me3 domains or both. Expression for genes associated with all putative enhancers located within introns is shown for comparison. **(B)** Bar graph showing combined score for Enrichr cell-type enrichment for the 5 cell types with the highest scores for each category. **(C)** Listings of the top ten GO terms for biological processes for the indicated groups; erythroid-specific terms are in boldface type.

**Supplementary Figure 6:**
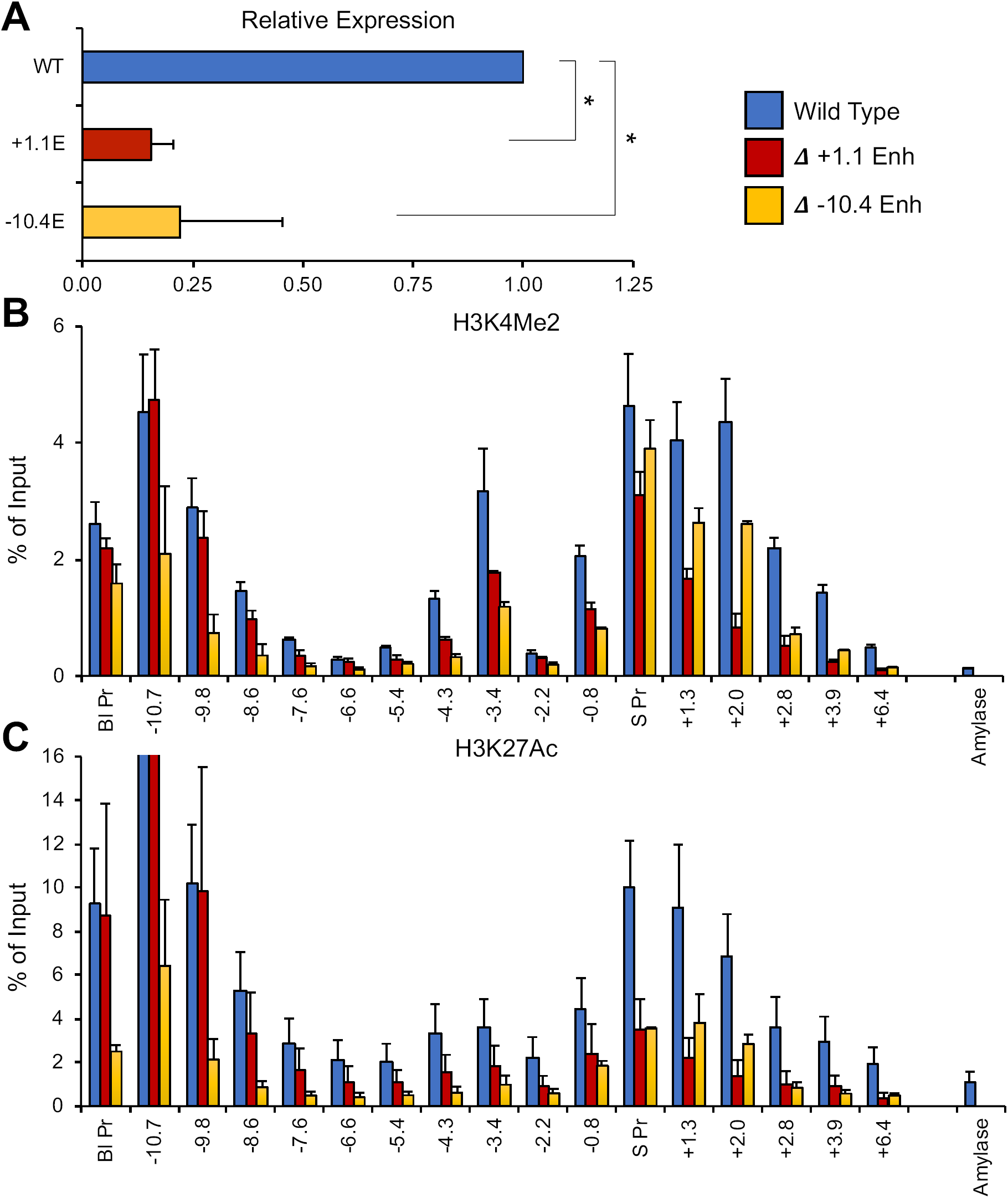
Effects of deletions of putative enhancers in the Band3 gene locus in differentiating MEL cells. **(A)** Normalized gene expression for Band3 measured by qrt-PCR in wt (MEL) and upon deletion of the indicated enhancer regions (+1.1 or −10.4). P-values were calculated using a two-tailed Student’s t-test;* = P<0.05. (B+C) Bar graph showing percent of input control obtained using PCR probes at the indicated locations (in kb) relative to the transcription start site for the Band3 gene in ChIP assays using antibodies specific for H3K4Me2 **(B)** or H3K27Ac **(C)** “Amylase” indicates a control probe within the inactive amylase gene locus.

